# Characterization of Sex Differences in Ocular HSV-1 Infection and Herpes Stromal Keratitis Pathogenesis of Wild Type and Herpesvirus Entry Mediator Knockout Mice

**DOI:** 10.1101/536185

**Authors:** Rachel E Riccio, Seo J Park, Richard M Longnecker, Sarah J Kopp

## Abstract

Sex differences related to immune response and inflammation play a role in the susceptibility and pathogenesis of a variety of viral infections and disease (S. L. Klein, Bioessays 34:1050-9, 2012). Herpes simplex virus type 1 (HSV-1) causes chronic inflammatory disease in the cornea, an immune privileged tissue, resulting in irreversible damage and blindness in affected individuals (A. Rowe, A. St Leger, S. Jeon, D. K. Dhaliwal, J. E. Knickelbein, and E. Vilain, Prog Retin Eye Res 32:88-101, 2013). Our research focuses on the role of Herpes Viral Entry Mediator (HVEM) as an immune regulator during ocular HSV-1 infection. Mice lacking HVEM (HVEM KO) exhibit lower immune cell infiltrates and less severe ocular disease in the cornea compared to wild type (WT) mice. As sex differences contribute to pathogenesis in many inflammatory diseases, we tested sex as a biological variable in the immune response to HSV-1 infection and Herpes Stromal Keratitis pathogenesis (HSK). Adult male and female WT and HVEM KO mice were inoculated with HSV-1 via corneal scarification and monitored daily for disease course. Viral titers and immune cell infiltrates were collected and analyzed. Our results indicate no significant difference in viral titers in tear film or affected tissues; immune cell infiltration; or clinical symptoms between males and females of either genotype. These results suggest that sex is not a significant biological variable in this experimental model, and that male and female mice can similarly be used in studies of ocular HSV-1 pathogenesis.

**IMPORTANCE:** Sex hormones have only recently been considered as an important factor for the development of certain diseases and as such should continue to be considered as a biological variable. Ocular Herpesvirus Type 1 (HSV-1), and the resulting Herpes Stromal Keratitis, is the leading cause of infectious blindness worldwide. We compared ocular HSV-1 infection and pathogenesis between sexes and found no significance difference between male and female wild type mice or herpesvirus entry mediator knockout mice. Therefore, male and female mice can be used interchangeably in studying ocular HSV-1 pathogenesis.

## INTRODUCTION

Sex differences play a role in the pathogenesis of a variety of viral infections including Influenza A, Epstein Barr, Hepatitis B and C, and Herpes simplex viruses (HSV) (1, 2). In these studies, differences in susceptibility and severity of the viral infections between males and females have been attributed to sex hormones which have roles in promoting immune cell function and cytokine synthesis (1–4). Notably, an epidemiological study of HSV type 2 (HSV-2) infections showed that females have a higher acquisition rate, incidence of symptoms, and prevalence of infection than males in genital infections (2, 5, 6). Whereas women have a greater risk of HSV-2 due to biologic susceptibility, men have a greater recurrence rate than women (5, 7). However, the rates and recurrence of HSV type 1 (HSV-1) infections are similar in men and women in genital infection (5). In a murine model of resistance to HSV-1, researchers found a sex-based difference in resistance in mice with the Herpes Resistance Locus, Hrl, which is a part of the tumor necrosis factor (TNF) superfamily, when challenged via ocular scarification with HSV-1: 52% of males were resistant to HSV-1 induced mortality, compared to 68% female mice (3). Although a clear sex bias has been shown in HSV-2 infections, differences in HSV-1 pathogenesis between males and females remains unclear.

Our laboratory studies herpesvirus stromal keratitis (HSK) which is an ocular disease that is the leading cause of infectious blindness worldwide (8, 9). It is typically caused by HSV-1 (10, 11). In our current study, we investigated if sex plays a role in this disease using a murine model. HSV-1 is a large double stranded DNA virus that enters and replicates in host cells mucosal membranes (12, 13). The virus then travels from the mucosal surfaces to sensory neurons, establishes latency, and persists in a latent state for the lifetime of the host. Reactivation can occur at any time with replication of virus traveling back to the mucosal surfaces. During ocular infection, reactivation of the virus from latency in the trigeminal ganglion to the corneal epithelium causes an inflammatory response characterized by corneal opacity and neovascularization which can ultimately lead to blindness due to these periodic episodes of virus reactivation (13–15).

Our previous studies have shown that Herpesvirus Entry Mediator (HVEM), a member of the TNF superfamily, has immunomodulatory functions and promotes ocular HSV-1 pathogenesis independent of viral entry receptor functions (16, 17). HVEM is a key immune regulatory protein that is part of the HVEM/LIGHT/BTLA/CD160 cosignaling pathway that regulates T cell activation and function (7). HVEM has a role in the adaptive immune responses of murine corneal inflammation and nerve damage that occur following ocular challenge of HSV-1 which is critical for the development of HSK (8). Mice lacking HVEM (HVEM KO) exhibit lower immune cell infiltrates and less severe ocular disease in the cornea compared to C57BL/6 wild type (WT) mice (8). Since a clear sex difference in mortality was observed in ocular studies with Hrl, we tested whether sex factors into our model of HSV-1 ocular infection. Moreover, sex hormones have been found to affect the expression of host cell surface proteins that function as receptors for viral entry such as the chemokine receptors for HIV-1, or DAF, a receptor for coxsackievirus. In these cases, estrogen promotes increased expression of receptors on B cells, which correlates with increased susceptibility of cells to infection and subsequent replication (2, 18, 19). In addition, immune response to pathogen infection can also vary between sexes. Most pertinent to our studies are those that show changes in cytokine expression which alters the immune response when males are compared to females. For example, when peripheral blood mononuclear cells from healthy men and women were stimulated with HSV-1 or Toll-like receptor 9 (TLR9) ligands, Interleukin 10 (IL10) production was significantly related with plasma levels of sex hormones in both groups (12). Males produced higher levels of IL10, which serves as a negative regulator of the response of both innate and adaptive immune cells, when stimulated with HSV-1 or TLR9 ligands compared to females (12). Previously, we used only male mice in our studies for characterizing HSK pathogenesis. Since HVEM modulates the immune response and sex plays a role in immune response in various viral infections, we specifically investigated if sex is a variable in infection by HSV-1 using WT mice or in the resulting immune response regulated by HVEM in HSK pathogenesis using HVEM KO mice. Moreover, little is known in regard to the role of sex in ocular health and disease (2, 10, 12).

Previous studies exploring the role of sex using different murine models of ocular HSV-1 have been contradictory. The experimental models have used other routes of infection, virus strains, and age at infection. In murine models of ocular HSV-1 infection, sex has been shown to play a role in only certain mouse backgrounds (14, 20). Female mice of the 129/Sv/Ev background display lower clinical periocular disease scores than both males and testosterone treated females although viral replication and corneal disease was similar, suggesting sex hormones enhance severity of clinical symptoms of HSK (20). In contrast, both male and female mice of the NIH/OLA background displayed similar clinical scores and viral titer from the trigeminal ganglia indicating sex does not play a role in pathogenesis of HSK in NIH/OLA background (21). Studies have also found sex differences in response to the HSV-1 glycoprotein D vaccine being more effective in women than men due to differences in immune cell response, where the gD peptide epitopes produced a higher CD4+ T cell response in females compared to males (22). Exploring the relationship between sex hormones and viral infection may offer insights into treatment and vaccination and a better understanding of the differences in immune cell response of males and females that may lead to novel therapies for human autoimmune diseases. Autoimmune diseases are more prevalent in women compared to men (1, 2). Eight percent of the population is affected by autoimmune diseases, where over two-thirds of those affected are woman. The stronger innate and adaptive immune response in females contributes to their increased susceptibility to inflammatory disease, and understanding whether this occurs in ocular HSV-1 infection and pathogenesis may suggest treatments that lessen severe disease humans (23).

In our current studies, we investigated sex specific differences in HSK pathogenesis in the C57BL/6 mouse strain between male and female mice to determine if sex plays a role in the inflammatory disease induced by the HVEM/LIGHT/BTLA/CD160 cosignaling pathway that regulates T cell activation and function (7). Our results indicate no significant difference between male and female mice when evaluating HSK disease pathogenesis. Our data is informative in determining the role of sex in this disease and allowing the use of both male and female mice in our ongoing studies and future experiments using the C57BL/6 mouse strain, which is commonly used for genetic knockout mouse lines.

## RESULTS

### HSV-1 Replicates to Similar Levels in Male and Female WT and HVEM KO Mice after Corneal Inoculation

In a murine model of infection, primary infection of the corneal epithelium results in detectable replication of HSV-1 for up to 6 days post infection when monitored in ocular tear film (11, 15). To determine whether sex impacts the replication of HSV-1, age-matched male and female adult WT and HVEM KO mice were inoculated with HSV-1 strain 17 via corneal scarification and viral titers of eye swabs of tear film were collected at days 1, 3, and 5 post infection. The titers showed no significant difference between male and female mice of either the WT or HVEM KO genotype at all 3 time points (Figure 1 A, C). Female WT mice had slightly higher titers than WT males on 3 dpi, and slightly lower titers than males on 5 dpi (Figure 1 A). Female HVEM KO mice had slightly lower eye swab titers than male HVEM KO mice on all three days (Figure 1 C). In all cases, these differences were not statistically significant.

**Figure 1.**
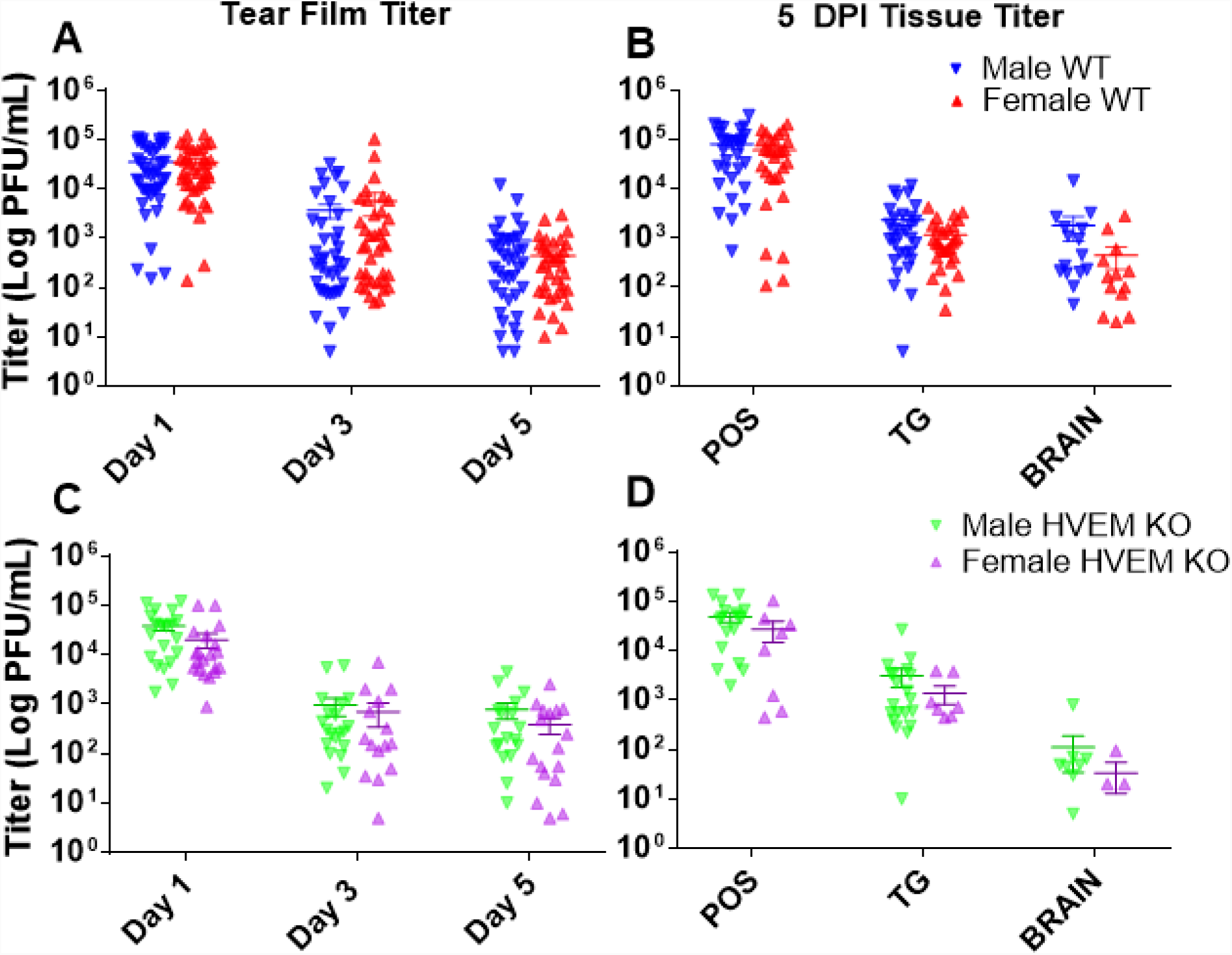
Male and female WT and HVEM KO mice exhibit similar HSV-1 eye swabs and tissues titers after corneal inoculation. Eye swabs were collected at days 1, 3 and 5 dpi and periocular skin (POS), trigeminal ganglia (TG), and brain tissues were collected at 5 dpi. (A) Titers from eye swabs of male and female WT mice (n=5-10 mice per group, 3 replicates). (B) Titers from tissues of male and female WT mice at 5 dpi (n=4-5 mice per group, 3 replicates). (C) Titers from eye swabs of male and female HVEM KO mice (n=5-10 mice per group, 3 replicates) and (D) from tissues of male and female HVEM KO mice (n=3-5 mice per group, 2 replicates) No difference was observed between either sex in each genotype. Titers calculated as means ± SEM were analyzed using two tailed t-tests with Holm-Sidak’s correction for multiple comparisons.

Following ocular infection, active replicating virus can also be quantified in the brain and surrounding tissues during acute phase of infection since it disseminates by retrograde transport to the neuronal cell bodies in the trigeminal ganglia (8, 11). To monitor dissemination, we collected periocular skin, trigeminal ganglia, and brain tissues at 5 days post infection to study this phase of the HSV-1 infection cycle (Figure 1 B, D). Although there was no significant difference between tissue titer in male and female mice within either genotype, lower titers were observed in tissues collected from female mice overall, but the differences were not significant when compared to male mice. From this data, we determine that sex has no significant effect on HSV-1 viral replication in the cornea, periocular skin, trigeminal ganglia, or brain of WT and HVEM KO mice.

### Male and Female WT and HVEM KO Mice have Similar Cellular Corneal Immune infiltrates during Acute and Chronic Inflammatory Phases of infection

We next investigated the corneal cellular immune response in WT and HVEM KO mice to determine if there were any sex differences in the prominent infiltration of inflammatory cells that occurs in stromal tissues following HSV-1 infection of the cornea. Previously, we found that cytokine responses in HVEM KO male mice were different when compared to WT male mice (7, 8). The early response to infection while virus is still actively replicating consists of polymorphonuclear leukocytes (PMN), mainly neutrophils, and is known as the acute inflammatory phase of infection. PMN infiltration peaks at around 2 dpi, declining at 5 dpi rising again during the chronic inflammatory phase of infection starting around 7 dpi after the virus has been cleared. Macrophages, dendritic cells, natural killer cells are also detected during this time and T cells enter the cornea during the chronic phase at 8-9 dpi (11). HSK is caused by this immunopathogenic response to infection and differences in infiltrating immune cell populations affect the overall severity of symptoms in the cornea (11, 15). An innate difference in cell-mediated immune responses exists between males and females, including response to viral infection. The reliance on subsets of CD4 helper T cells differ between males and females in overcoming infection, where females have been reported to have higher Th1 and IFN-y responses than males (2, 24, 25). Androgens have been found to enhance viral and immune factors, where estrogens modulate both innate and adaptive immunity to be protective (26).

Cornea pairs of infected male and female WT and HVEM KO mice were collected during the acute phase of infection at 5 dpi (Figure 2 A, B) and then at the chronic inflammatory phase of infection at 14 dpi (Figure 2 C, D) to determine if the immune cell populations infiltrating the corneas of male and female adult mice differed during either phase of infection. Cell populations were measured via flow cytometry as described in the materials and methods by first gating on populations of CD45+. The count of infiltrating cells from CD45+/CD11b+ myeloid lineages during the acute inflammatory phase of infection were measured (Figure 2 A). These populations include Ly6G-/CD11c+ myeloid dendritic cells (mDCs), Ly6G-/CD11c- monocyte and macrophages (M), Ly6G-/CD11c-/Ly6C+ inflammatory macrophages (I-M), Ly6G-/CD11c-/Ly6C- non-inflammatory macrophages (I-M), and Ly6G+/CD11c+ polymorphonuclear neutrophils (PMN). The count of infiltrating cells from CD45+/CD3+ lymphoid lineages during the acute phase were also measured (Figure 2 B). These populations include CD4+CD8-CD4 t cells (CD4+), CD4-CD8+ CD8 t cells (CD8+), NK1.1+ natural killer t cells (NKT), and CD4-CD8-NK1.1-double-negative T cells (DNT). The myeloid and lymphoid cell populations during chronic inflammatory phase were also measured (Figure 2 C, D). We analyzed these specific populations, including the CD4+ T cells and PMN neutrophils since they contribute to most of the damage to the eye during HSK (11). We found, as expected, the WT male mice had a significantly higher number of PMN cells and an increase in number of CD4+ T cells during chronic phase of infection than the HVEM KO male mice (8). This difference between genotypes was also observed in the female mice. Overall, male and female mice exhibited similar levels of immune cell infiltration in each genotype. The data suggests sex does not affect immune cell infiltrates in the cornea during HSV-1 ocular infection.

**Figure 2.**
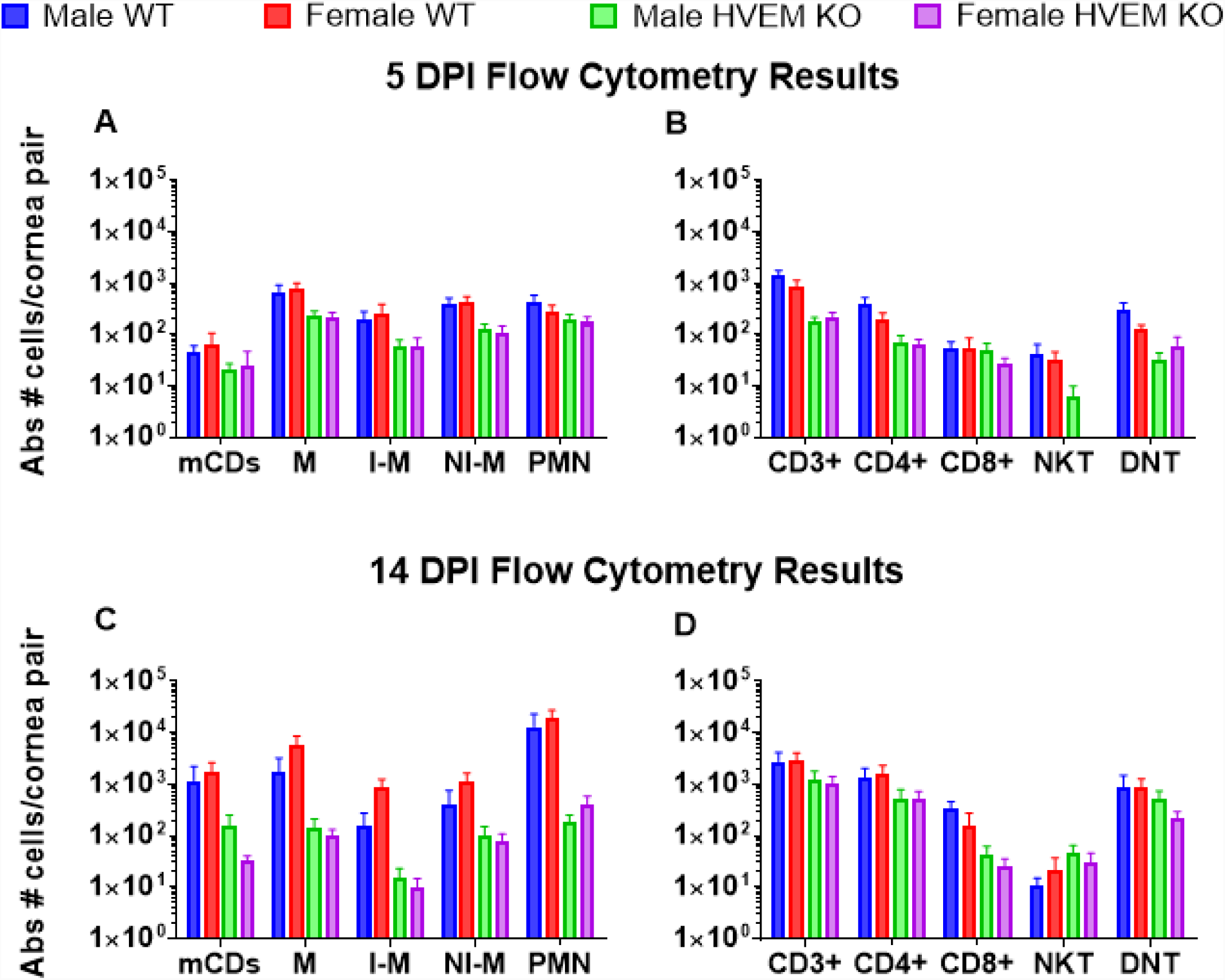
Immune cells infiltrate male and female WT and HVEM KO corneas at similar levels during acute (5dpi) and chronic (14dpi) phases of HSV-1 corneal infection. (A & B) Flow cytometry analysis of immune cell infiltrates in WT and HVEM KO corneas at 5 dpi (n = 8-10, two replicates) and (C & D) 14 dpi. (A) Absolute number of myeloid infiltrates of dendritic cells (mDCs), macrophages (M), inflammatory macrophages (I-M), non-inflammatory macrophages (NI-M), and polymorphonuclear leukocytes (PMN) at 5 dpi and (C) at 14 dpi. (B) Absolute number of lymphoid infiltrates of CD4+ T cells, CD8+ T cells, NK T cells, or DN T cells per cornea pair at 5 dpi and (D) at 14 dpi. (n=8-20 mice, 2-3 replicates). There was no significant difference between males and females of either genotype. The difference between WT and HVEM KO is consistent with our previous studies. Infiltrating cell percentages calculated as means ± SEM were analyzed using two tailed t-tests with Holm-Sidak’s correction for multiple comparisons.

**Figure 3.**
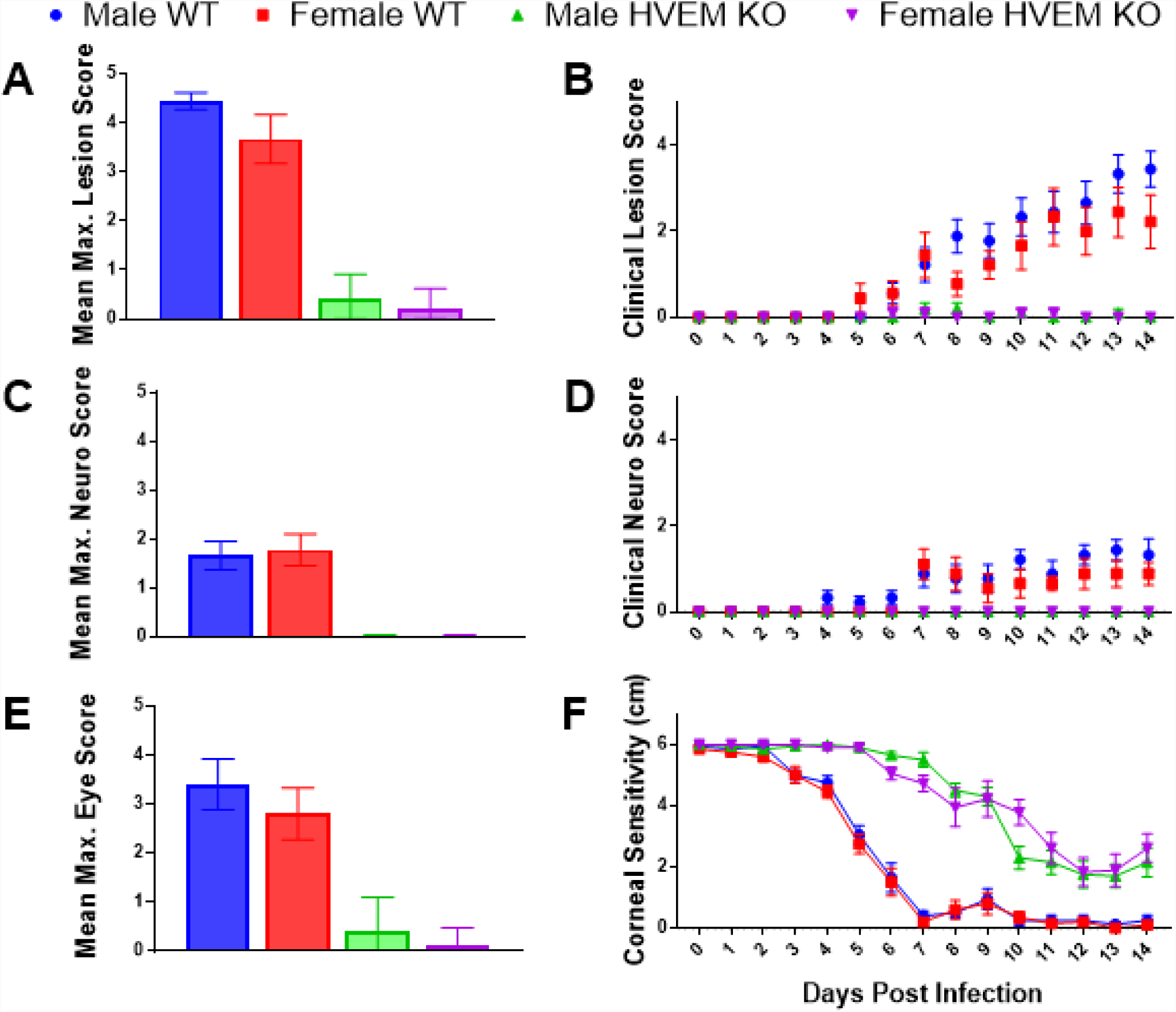
Male and female mice exhibit similar clinical symptoms during chronic inflammatory phase of HSV-1 corneal infection. Clinical symptoms of each mouse were scored and recorded each day from inoculation to chronic inflammatory phase, day 0-14. Mice were scored on severity and presence of lesions (A and B), neurological disease (C and D), and eye disease (E), as well as corneal sensitivity (F) and weight loss (not shown). (A) Maximum lesion score, (C) maximum neurological disease score, and (E) maximum eye disease score within 14 dpi. (B) Lesion score and (D) neurological disease score from 0 to 14 dpi. (F) Corneal sensitivity from 0 to 14 dpi. (n=5 mice per group, 2 replicates). No difference was observed between the sexes. Scores and sensitivity were calculated as means ± SEM were analyzed using two tailed t-tests with Holm-Sidak’s correction for multiple comparisons.

### Clinical Symptoms of Corneal HSV-1 Pathogenesis is Similar in Male and Female WT and HVEM KO Mice

Mice infected with HSV-1 not only exhibit ocular disease but neurological symptoms including seizures, loss of balance, and weight loss. To determine whether sex plays a role in clinical symptoms of HSK and pathogenesis of corneal infection, male and female WT and HVEM KO were scored for clinical symptoms daily from 0 to 14 dpi (Figure 4). As described in Table 1, scores were based on appearance and severity of lesions, neurological symptoms, and clinical eye disease on a scale of 0-5 with 0 being not present and 5 being most severe. The mean maximum lesion score was 4.4 for male WT mice and 3.7 for female WT mice; 0.4 for male HVEM KO mice, and 0.2 for female HVEM KO (Figure 4 A). The severity of lesions is similar between males and females within each genotype. The occurrence of lesions and mean score on each day was also measured (Figure 4 B). Male and female HVEM KO mice experienced similar progression of lesions, and female WT mice had slightly lower scores than male WT mice. The neurological scores of male and female mice was recorded and the mean maximum score was calculated. The mean maximum neurological score was 1.7 for male WT mice and 1.8 for female WT mice; HVEM KO mice did not display any neurological symptoms as we have previously seen (Figure 4 C). The severity of neurological symptoms is similar between males and females within each genotype. The onset of neurological symptoms and mean score on each day were measured (Figure 4 D). Male and female mice experience similar levels of neurological symptoms throughout HSV-1 pathogenesis. Additionally, eye disease score of male and female mice were taken. The mean maximum eye disease score was 3.4 for male WT mice and 2.8 for female WT mice; 0.4 for male HVEM KO mice, and 0.13 for female HVEM KO mice (Figure 4 E). The severity of eye disease symptoms is also similar between males and females within each genotype. Overall, female mice exhibited attenuated clinical symptoms compared to male mice in both genotypes, but these differences were not significant. Our results with female and male mice were identical to our previous studies with male mice showing that HVEM contributes to more severe disease in WT mice compared to HVEM KO mice (16).

**Table 1:** Clinical Symptom Scoring Scale for HSK in Mice.

| Lesion Score |  | Neurological Score |  | Eye Score |  |
| --- | --- | --- | --- | --- | --- |
| 0 | Lesion Free | 0 | Symptom Free | 0 | Symptom Free |
| 1 | Small area of broken skin <0.5cm | 1 | Ruffled fur, hunched posture, normal movement | 1 | Pus around edges of eyes |
| 2 | Area of broken skin 0.5-1cm | 2 | Hunched posture, slow movement | 2 | Pus and squint |
| 3 | Broken skin, bleeding, scabbing, or pustules | 3 | Hunched posture, slow movement, labored breathing | 3 | Closed |
| 4 | Broken skin >1cm with multiple pustules or scabbing | 4 | Hunched, labored breathing, little to no movement | 4 | Closed, scab formation |
| 5 | Severe scabbing or bleeding with pustules | 5 | Moribund or dead | 5 | Severe scabbing |

A major symptom of HSK includes corneal scarring which leads to loss of corneal sensitivity and vision. Esthesiometry, which measures corneal sensitivity is based on the shortest filament length required to elicit a blink response, was used to monitor corneal sensitivity from 0 to 14 dpi. With a 6cm length filament representing no loss of sensitivity and 0.5cm or less representing total loss of sensitivity (Figure 4 F). Male and female mice had similar corneal sensitivity throughout corneal infection. Our previous data showed that HVEM contributes to loss of corneal sensitivity by comparing HVEM KO to WT adult male mice, where WT mice lose sensitivity and HVEM KO mice retain sensitivity (8). The same result is observed in female mice. From this we conclude sex does not play a significant role in the clinical disease course during HSV-1 ocular infection of mice.

## DISCUSSION

Susceptibility to viral infections is often reduced in females due to a greater humoral and immune response to vaccination and infection when compared to males (2). Historically, only one sex was used during murine models of HSV-1 infection, and male-predominant populations have been used in human studies and trials (2, 10). Studies have shown sex dependence of HSV infection of the central nervous system where females were more susceptible than males (27, 28). Female mice have also been shown to have higher HSV titers from brain tissue when inoculated via intraperitoneal injection, resulting in greater mortality and severity of disease than males (1, 28). Estrogen treatments, castration, or hormones do not fully explain these sex differences (1, 20). In human studies, the high prevalence of HSV-1 in the trigeminal ganglia (TG) was shown not to be related to sex, with 88% of females and 90% males having HSV-1 positive TG (29). These studies reveal the importance of addressing and understanding differences in susceptibility to HSV-1 infection and ocular disease between the sexes since it may result in more effective therapies and vaccines.

Our results indicate viral titers in tear film and tissues from adult HVEM KO and WT mice tissue were not affected by sex (Figure 1). The data revealed a trend in female mice to have slightly lower tissue titers than male mice of both the WT and HVEM KO genotypes. These findings are similar to those of an ocular model using the 129-mouse strain background, in which viral titer in the TG was slightly lower in females but not significant (20). In contrast, HSV-2 infection does show a sex bias in human studies in which women exhibit a higher prevalence of infections than males in all age groups, and seronegative women acquire virus faster and have a higher incidence of symptomatic infections than males (2, 6, 30). Our previous research showed HVEM is not required for disease during HSV-2 infection of mice, however, HVEM is required for full HSV-1 pathogenesis of in eye following infection (8). Our data suggests rather than being influenced by sex hormones as in HSV-2 infection, the immune response in HSV-1 pathogenesis is being modulated by HVEM. This may suggest why sex differences have been reported in HSV-2 but have not been shown in our research of ocular HSV-1 pathogenesis. Another study found some effects of the interferon gamma (IFN-y) receptor to be sex specific in ocular HSV-1 infection. In IFN-y receptor knockout mice, females displayed less severe POS disease scores than males, and testosterone-induced females produced male-like POS symptoms (20). However, the effect of sex hormones on host receptors like HVEM has yet to be fully understood, and further investigation into what is contributing to the reduced infiltrates and less severe disease in mice lacking HVEM is needed.

Sex hormones may not influence the immune response in HSK because of the immune privilege of the cornea which is the site of infection. Previous studies have shown the development of corneal eye disease to be strongly correlated with an inflammatory response characterized by CD4+ Th1 cytokine-producing T cells (20, 31). The results shown in Figure 2 analyzing CD4+ cells found males had a higher number of CD4+ cells in response to infection during the acute stage whereas females had a slightly higher CD4+ response during chronic infection. In the chronic inflammatory stage of infection which contributes to the severity of HSK, no significant difference was observed in immune cell infiltrates (Figure 2 C, D). HVEM is known to influence HSV pathogenesis in the ocular murine model during most stages of infection, from entry and acute viral replication, innate responses, chronic inflammation, and in viral latency (8, 32–35). Our previous findings reported that HVEM contributes significantly to increased immune cell infiltrates during ocular HSV-1 infection (17). Since no significant differences were observed between males and females within each genotype, and the differences between the genotypes was consistent within the sexes, we conclude sex does not play a role in the innate or adaptive response to ocular HSV-1 infection.

In addition, the difference in clinical symptoms between WT and HVEM KO that we observed in our previous studies confirm that HVEM modulates the immune response rather than being altered by sex (17). Our results also show no significant differences in clinical symptoms between males and females of either genotype in our experimental model. Female mice trended toward slightly lower clinical scores than male mice in both the WT and HVEM KO genotypes (Figure 4). A previous study using the HSV-1 KOS strain found that males displayed less severe clinical symptoms (disease score mean of 1.5) than females (score mean of 2.0) on average (14). This was also observed in another study in which clinical eye disease scores were slightly lower in females but not significant (20). Clinical examination using a slit-lamp microscope may provide better diagnosis and scoring of HSK as this the primary diagnostic tool in patients (36, 37). However, there were no sex differences in corneal eye disease scores in our results and the other reports of ocular HSV-1 infection and clinical HSK pathogenesis (14, 20).

Our results suggest sex is not a significant biological variable in ocular HSK in the C57BL/6 mouse background and does not contribute to the observed reduction of immune cell infiltration and less severe clinical symptoms in HVEM KO mice. Overall, we determined there was no significant difference in HSV-1 ocular infection in male and female mice. In compliance with the NIH standards of considering sex a biological variable, we report that both male and female mice can be used simultaneously in studies using C57BL/6 mice and HVEM KO mice made on the C57BL/6 background.

## MATERIALS AND METHODS

### Ethics Statement

All experiments utilizing mice were performed in strict adherence to the recommendations in the Guide for the Care and Use of Laboratory Animals of the National Institutes of Health. The Institutional Animal Care and Use Committee of Northwestern University approved the protocol (Protocol no.IS00001532). Procedures were performed under anesthesia using a ketamine/xylazine cocktail or under isoflurane anesthesia. Minimization of suffering was prioritized.

### Cells and Viruses

African green monkey kidney cells (Vero) were used for propagation of virus and all viral plaque assays. HSV-1 strain 17 was obtained from David Leib (Dartmouth Medical School, Hanover, NH)

### Viral Plaque Assay

Viral plaque assay using Vero Cells was performed to determine viral titers of swabs and tissues, as previously described (17).

### Animals and Procedures

Mice were maintained in a specific-pathogen-free environment and were transferred to a containment facility after infection. 8-15 week old male and female C57BL/6 mice were used for all experiments. Mice were bred in house. Wild type mice were acquired from Jackson Laboratory (WT mice) and *Tnfrsf14−/−* mice (HVEM KO mice) were obtained from Yang-Xin Fu (38).

For ocular inoculation, mice were anesthetized with an intraperitoneal injection of ketamine/ xylazine solution. Corneas were abraded lightly with a 25 gauge needle in a crosshatch pattern and 5ul of 2 x 10^6 PFU HSV-1 strain 17 was applied to each cornea. Mice were weighed and scored for clinical symptoms daily. Clinical scores were divided into observing lesions and neurological symptoms. The scoring scale, used in our previous studies, is shown in Table 1.

Mice displaying a neurological score of 4 or weight loss greater or equal to 30% starting weight at infection were sacrificed and scored as 5. Eye swabs were collected after mild anesthesia with isoflurane where mice were unresponsive to footpad prick. The eye was gently proptosed and a sterile cotton swab pre-moistened with DMER was wiped three times around the circumference of the eye and twice across the center of the cornea in an “X” shape as previously described (16). Swabs were then placed into 1 mL of DMER media (DMEM containing 5% (vol/vol) FBS, 1% gentamicin, 1% ciprofloxacin, and 1% amphotericin B) and stored at −80°C. To determine viral titers, samples were thawed and vigorously vortexed for 30 seconds prior to performing a plaque assay.

Two experimental groups of mice were used. Mice were sacrificed on day 5 for flow cytometry, eye swabs, and for harvest of tissues. The second group were monitored for clinical symptoms, eye swabs were taken for viral titers, and on day 14 mice were sacrificed for flow cytometry. For day 5 experiments animals were sacrificed and the POS, TG, and brains were collected as previously described (6, 7, 17). After dissection, all samples were placed in 1 mL DMER, homogenized, sonicated, and stored at −80°C until titration. Brain samples were centrifuged prior to titration to remove debris.

### Flow Cytometry

On either day 5 or day 14 post infection corneal pairs and spleens were collected from individual mice into cold Phosphate Buffered Solution (PBS). Corneas were placed into Liberase (Roche, Indianapolis, IN, USA) at 0.7mg/mL in RPMI media to digest for 1hr at 37°C, 5% CO2 incubator. Corneas were homogenized through a 100um mesh with a 1ml syringe plunger. Cells were washed with cold PBS, centrifuged at 4°C and then strained through a 40um mesh and collected into 300ul. Spleens were prepared in a similar manner as the corneas with an added red blood cell lysis incubation between the 100um and 40um straining steps. Spleens were collected into 3mL. Live cell counts were obtained using a Countess Cell Counter. All of the cornea cells and a portion of the spleen cells were each incubated at 1:1000 of Live/Dead Fixable Aqua Dead Cell Stain Kit (Thermo Fisher Scientific) in PBS in the dark at RT for 30 minutes. Samples were washed with PBS and incubated with Fc block (0.5–1.0ug/sample anti-mouse CD16/CD32 [eBioscience, San Diego, CA, USA] in PBS 1% fetal bovine serum 0.1% sodium azide [FACS buffer]) for 5 minutes at 4°C in the dark. Conjugated antibodies (2ug/mL final per sample) were added directly to Fc block, then incubated for 1 hour at 4°C in the dark.

The following antibodies were used: HVEM anti-mouse APC (HMHV-1B18), Ly6G Brilliant Violet 421(1A8), CD4 PE(RM4-5), CD8a Brilliant Violet 421 (53-6.7) from BioLegend; CD45 FITC (30-F11), CD3 APC eFluor 780 (17A2), Cd11b PE Cy7 (M1/70), Cd11c PE(N418), Ly6C PerCP Cy 5(HK1.4), CD3e PE Cy7 (145-2C11)from Ebioscience.; NK1.1 APC Cy7(PK136) from BD Biosciences. Isotype controls were all from Ebioscience: Arm Ham IgG isotype control APC (eBio 299 Arm) isotype, Rat IgG2bk isotype control PerCP eFluor 710 (eB149/10HS), Rat IgG2a isotype control APC eFluor 780 (eBR2a). Sample data was acquired using the FACS Canto II (BD Biosciences), the entire corneal pair sample was run, and the spleen sample run was stopped at 100,000 live cells. The FlowJo 10.1 software (Ashland, OR, USA) was used for data analysis. All samples were first gated on Lymphocytes. Single cells were separated according to FSCA vs FSCH. Live cells isolated using the Live/Dead plot. Cell populations were measured via flow cytometry gating on populations of CD45+ cells. CD45+/CD11b+ myeloid lineages were gated into further populations of Ly6G-/CD11c+ myeloid dendritic cells (mDCs), Ly6G-CD11c-monocyte and macrophages (M), Ly6G-/CD11c-/Ly6C+ inflammatory macrophages (I-M), Ly6G-/CD11c-/Ly6C- non-inflammatory macrophages (I-M), and Ly6G+/CD11c+ polymorphonuclear neutrophils (PMN). CD45+/CD3+ lymphoid lineages were gated into further populations of CD4+CD8-CD4 t cells (CD4+), CD4-CD8+ CD8 t cells (CD8+), NK1.1+ natural killer t cells (NKT), and CD4-CD8-NK1.1-double-negative T cells (DNT).

### Corneal Sensitivity

To determine sensitivity of the central cornea, a Luneau Cochet-Bonnet esthesiometer (No. WO-7760; Western Ophthalmics, Lynnwood, WA, USA) was used to measure the blink threshold. Animals were scruffed, and the length of the monofilament was varied from 6.0 to 0.5 cm and applied perpendicularly to the surface of the central cornea until the first inflection point. A positive response was recorded when two or more blinks were obtained out of three attempts. Mice with an absence of a blink response at 0.5 cm were scored as a 0. The same examiner performed all measurements.

### Statistics

Geometric means of numbers of viral-plaque-forming units per tissue sample, maximum neurologic and lesion scores, were compared using the unpaired Student’s t-test with Holm-Sidak’s multiple-comparison test. The unpaired Student’s t-test and Holm-Sidak’s multiple-comparison test were used to determine variance over time between groups with respect to development of lesions or neurologic morbidity. GraphPad Prism 7.0 software for statistical analysis.

## ACKNOWLEDGEMENTS

We are grateful for our funding source: National Eye Institute 1R01EY023977-01A1.

We would like to thank Nanette Susmarski for her expert assistance with cell culture, David Leib for providing the HSV-1 strain 17 which were used in our studies, and Yang-Xin Fu for the HVEM KO mice. We also thank fellow members of the Longnecker lab for their support.

